# Phospho-β-catenin expression in primary and metastatic melanomas and in tumor-free visceral tissues

**DOI:** 10.1101/2020.09.13.295410

**Authors:** Joel Pinczewski, Rebecca Obeng, Craig L. Slingluff, Victor H. Engelhard

**Affiliations:** Department of Pathology, University of Virginia School of Medicine, Charlottesville, VA, 22908; Department of Microbiology, Immunology and Cancer Biology, University of Virginia School of Medicine, Charlottesville, VA, 22908; Beirne Carter Center for Immunology Research, University of Virginia School of Medicine, Charlottesville, VA, 22908; Department of Surgery, University of Virginia School of Medicine, Charlottesville, VA, 22908; UVA Cancer Center, University of Virginia School of Medicine, Charlottesville, VA, 22908

## Abstract

β-catenin (βcat) is an important downstream effector in the Wnt signaling pathway and plays an important role in the development and progression of many cancers including melanoma. βcat expression is regulated by GSK-3β-mediated phosphorylation at positions 33, 37 and 41. In normal cells, phosphorylation at these sites triggers proteasomal degradation, which in turn prevents accumulation of free cytoplasmic βcat. In cancer cells, stabilized β-catenin translocates into the nucleus, where it associates with TCF/Lef proteins to activate transcription of genes that promote tumorigenesis and metastasis. It has been suggested that nuclear phospho-βcat (pβcat) staining may be diagnostically useful in differentiating primary from metastatic melanoma. Also, a pβcat peptide (residues 30-39, p33) is naturally presented by melanoma cells as a T-cell target. We evaluated the expression of pS33-βcat in primary and metastatic melanoma tissues by immunohistochemistry. pS33-βcat was detected in primary and metastatic melanomas and was most commonly cytoplasmic and almost never exclusively nuclear. Interestingly, staining with pS33-βcat and pS33/37/T41-βcat antibodies was most intense in mitotic melanoma cells, consistent with prior studies demonstrating changes in the level of βcat during cell division. We observed no significant differences in pβcat staining location or intensity between primary and metastatic melanomas, suggesting that pβcat may have limited diagnostic or prognostic utility in melanoma. However, the high expression in dividing cells suggests promise as an immunotherapeutic target.

## Introduction

β-catenin (βcat) forms a downstream portion of the highly conserved canonical Wnt cell signaling pathway. As a member of this pathway, βcat has a pivotal role in embryogenesis and in cell signaling in later life. Mutations in βcat or the proteins that regulate it (e.g., the adenomatous polyposis coli (APC) protein), which lead to accumulation of βcat, are associated with the development and progression of a variety of benign and malignant neoplasms, including familial adenomatous polyposis, Gardner’s syndrome, and colorectal cancer(1-4). However, mutations resulting in βcat accumulation are also associated with other malignancies including gastric and endometrial adenocarcinomas, hepatocellular cancers, hepatoblastomas, and melanomas(2, 5). In unstimulated adult cells, βcat is located mainly on the cytoplasmic side of the cell membrane, where it associates with E-cadherin as part of adherens junctions(6). Free cytoplasmic βcat is maintained at a low level by formation of a destruction complex involving βcat, glycogen synthase kinase-3 β (GSK-3β), casein kinase 1a (CK1a), APC and Axin(7). This leads to sequential phosphorylation of the complexed βcat at T41, S37, and S33 by GSK-3β, targeting it for destruction by the ubiquitin/proteasome pathway. Signaling through the Wnt pathway results in a different fate for βcat(8). In this setting, free βcat is protected from phosphorylation and is able to accumulate in the cytoplasm. This build-up in cytoplasmic βcat leads to its eventual translocation to the nucleus where it complexes with T-cell factor (TCF)/ lymphoid enhancer-binding factor (LEF) and other molecules and effects transcription of genes involved in tumorigenesis and metastasis such as c-myc and metalloprotease(9-11).

Zarling et al have identified a βcat peptide spanning amino acids 30-39 and phosphorylated at S33 on melanoma, which is presented by human HLA-A2 molecules as an antigen for cytotoxic T cells(12). This suggests that N-terminal phosphorylation of βcat occurs in melanoma cells and the resulting peptide may be a target for cancer immunotherapy. Phosphorylation at S33 only occurs after phosphorylation at residues 37 and 41; so, detection of pS33-βcat identifies the tri-phosphorylated form of βcat. pS33-βcat has not been shown to have transcriptional activity, but some authors have reported that the presence of cytoplasmic and nuclear pS33-βcat staining in malignant cells has diagnostic and prognostic significance. Nakopolou found that cytoplasmic pS33-βcat staining was associated with an improved outcome in breast cancer, whereas high levels of nuclear pS33-βcat staining were associated with a poor outcome(13). Kielhorn found that most primary and metastatic melanomas did not stain with an antibody detecting pS33-βcat(14). When present, however, pS33-βcat staining was always nuclear and was more commonly seen and more intense in metastatic melanoma than in primary melanomas. On the basis of this and Kaplan-Meier survival curves, Kielhorn suggested that nuclear pS33-βcat staining and its intensity might serve as a useful diagnostic and prognostic marker in melanoma(14).

Activation of the Wnt/βcat pathway in melanoma has been reported to have immunological significance, with reduction in immune cell infiltrates in melanomas with Wnt/βcat activation(15). This heightens interest in understanding the relevance of pS33-βcat expression in melanoma, coupled with the finding that a pS33-βcat peptide is an epitope recognized by human CD8^+^ T cells. The relevance of pS33-βcat as a T cell target requires understanding whether pS33-βcat is expressed in normal tissues from vital organs and whether it is selectively expressed by melanoma cells compared to such normal tissues. βcat is expressed in normal tissues including heart and lung, but regulation of βcat phosphorylation is complex enough that expression levels of βcat may not predict expression of pS33-βcat. To our knowledge, the expression of pS33-βcat in human normal tissues has not been assessed. We hypothesized that pS33-βcat would be expressed at higher levels in melanoma metastases than in normal tissues. Thus, in the present study, we performed immunohistochemistry (IHC) studies to determine the breadth of pS33-βcat expression in primary and metastatic melanoma tissues as well as in normal human tissues. We also compared the levels and intracellular locations of pS33-βcat staining in primary and metastatic melanoma specimens as well as pS33-βcat staining in normal tissues to re-evaluate the potential use of pS33-βcat as a diagnostic and prognostic indicator for melanoma.

## Material and Methods

Melanoma tissues for analysis in this study included samples of formalin-fixed and paraffin-embedded (FFPE) specimens from a melanoma tissue microarray (TMA), preparation of which has been described previously(16), plus additional FFPE samples obtained from the archives of the Department of Anatomic Pathology at the University of Virginia (IRB-HSR 15237 and 10598). These included fifteen primary melanomas and forty-three metastatic melanomas. Included among the metastatic melanoma specimens were lung (n=5), heart (n=1) and liver (n=1) specimens that contained melanoma metastases and evaluable portions of non-malignant (“normal”) tissue from the same organs that had been invaded. Cells of the human melanoma cell line SLM2(17) growing in log phase also were pelleted, formalin fixed, paraffin embedded, and then sectioned for the study. Tissue sections were deparaffinized in xylene and rehydrated by sequential passage through graded alcohol/water solutions. Heat-induced antigen retrieval was performed using Vector’s Target Retrieval Solution (Vector Laboratories, Burlingame, CA), pH 6 at 100°C for 20 min.

Working concentrations of antibodies to βcat (1:500, Epitomics, Burlingame, CA # 1247-1) and phospho-βcat (pS33-βcat 1:200, Santa Cruz, Santa Cruz, CA # SC-16743-R; pS33/37/T41-βcat 1:1200, Cell Signaling Technology, Danvers, MA, #9561) were used. In some assays, a 13-mer pS33-βcat blocking phosphopeptide (QSYLDpSGIHSGAT, Genscript, Piscataway, NJ) was added to anti-βcat and anti-pS33-βcat antibodies (1:200), giving a final peptide concentration of 50 μg/ml, and incubated for 30 min prior to being added to slides for blocking. The slides stained for pS33-βcat were incubated overnight at 4 degrees, and slides stained for βcat were incubated for 30 min at room temperature. Expression was detected using Vector’s ImmPRESS peroxidase kit and visualized with Vector’s ImmPACT(tm) AEC substrate. Slides were then counterstained with Gill’s hematoxylin. Negative control slides were prepared by omitting the primary antibodies. Slides were scanned using an Aperio CS slide scanner (Aperio, Vista, CA) and were analyzed using the Aperio Imagescope “positive pixel count” analytical software algorithm. The resulting data were used to calculate total staining density per unit area.

## Results

The specificity of the polyclonal pS33-βcat antibody was tested by staining four normal tissues -- liver, placenta, spleen and kidney – with and without pre-incubation with a corresponding pS33-βcat phosphopeptide as a control. As a further control, a commercial anti-βcat antibody was also used. All four normal tissues stained strongly with both the pS33-βcat and βcat specific antibodies (**Figure 1**). Staining with the pS33-βcat antibody was significantly reduced by pre-incubation with pS33-βcat phosphopeptide. Pre-incubation with the blocking phosphopeptide had minimal effect on βcat antibody staining.

**Figure 1.**
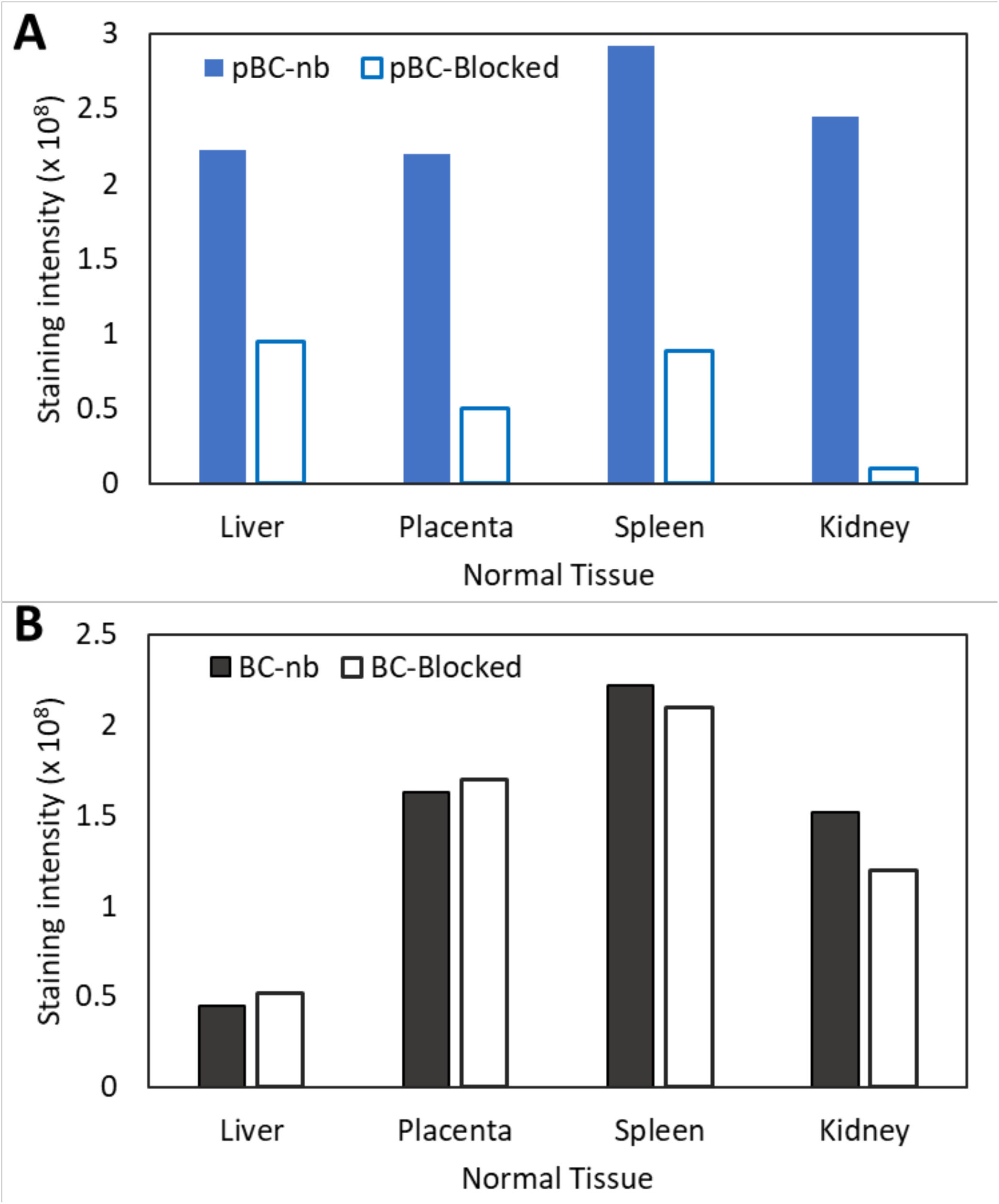
Phospho-β catenin in normal human tissues. Total staining density per unit area for normal liver, lung, spleen and placenta stained with antibodies to (**A**) pS33-βcat and (**B**) βcat, with pre-incubation with pS33-βcat blocking phosphopeptide (pβcat-Block and βcat-Block) or without blocking (pβcat-nb and βcat-nb).

To assess the diagnostic and prognostic value of pS33-βcat, we compared the intensity and staining location of the pS33-βcat antibody in TMA blocks containing 15 primary and 43 metastatic melanomas. All fifteen primary melanomas (100%) had cytoplasmic staining (**Figure 2**). In addition, 2/15 (15%) had combined membranous and cytoplasmic staining and 3/15 (20%) had combined nuclear and cytoplasmic staining. In the single remaining case, membranous, cytoplasmic, and nuclear staining were all evident concurrently. Among the metastatic melanomas, 36/43 (84%) had cytoplasmic staining. Nuclear staining was observed in 6/43 cases (14%). However, in all but one of these cases the nuclear staining was combined with cytoplasmic staining. Therefore, isolated nuclear staining was only observed in one case (2%). Membranous staining was evident in 5/43 cases (11%). These consisted of 4 tumors with combined nuclear and cytoplasmic staining and 1 with isolated membranous staining. A complete lack of staining was observed in 7/43 cases (16%).

**Figure 2.**
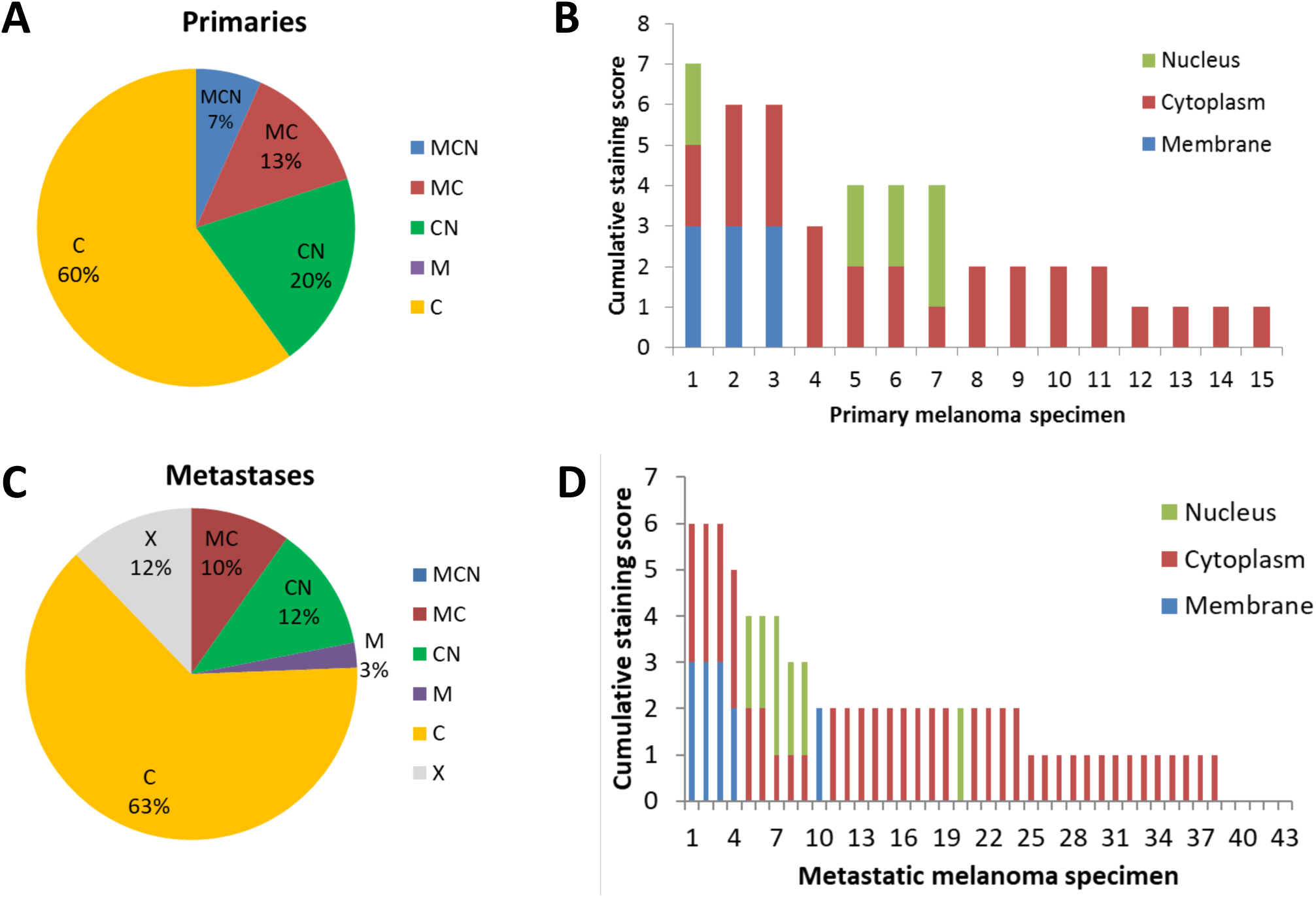
Expression and localization of phospho-β-catenin in primary (**A**, **B**) and metastatic (**C**, **D**) melanomas. (**A, C)** The proportion with nuclear (N), cytoplasmic (C), membranous (M) staining, combinations (MCN, MC, CN), or no staining (X). (**B, D**) The extent of staining in each region is shown per patient tumor based on scoring of 0 to 3 in each cellular area (N, C, and M). The intensity of staining was graded from 0 (no staining) to 3+ (heavy staining).

To explore whether there were differences in the level of pβcat in tumors and normal tissues that might have relevance in the diagnosis and treatment strategies for melanoma patients, we compared staining in melanoma metastases versus adjacent lung, liver, and heart tissue that did not contain melanoma. Whole sections of lung, liver and heart tissue, containing both metastatic melanoma and adjacent uninvolved tissue, were stained with the pS33-βcat antibody. Metastatic melanoma in the lung tissue mainly had cytoplasmic staining but also included one case with minimal staining and another with predominantly membranous staining (**Figure 3**). The most intense staining occurred in the uninvolved lung tissue and in respiratory epithelia. As shown in **Figure 4**, cardiac myocytes had higher staining intensities than the metastatic melanoma in heart tissue, with the most intense staining occurring in the intercalated discs. In liver tissue (**Figure 5**), there was only very weak staining for both metastatic melanoma and adjacent uninvolved hepatocytes (**Figure 5**).

**Figure 3.**
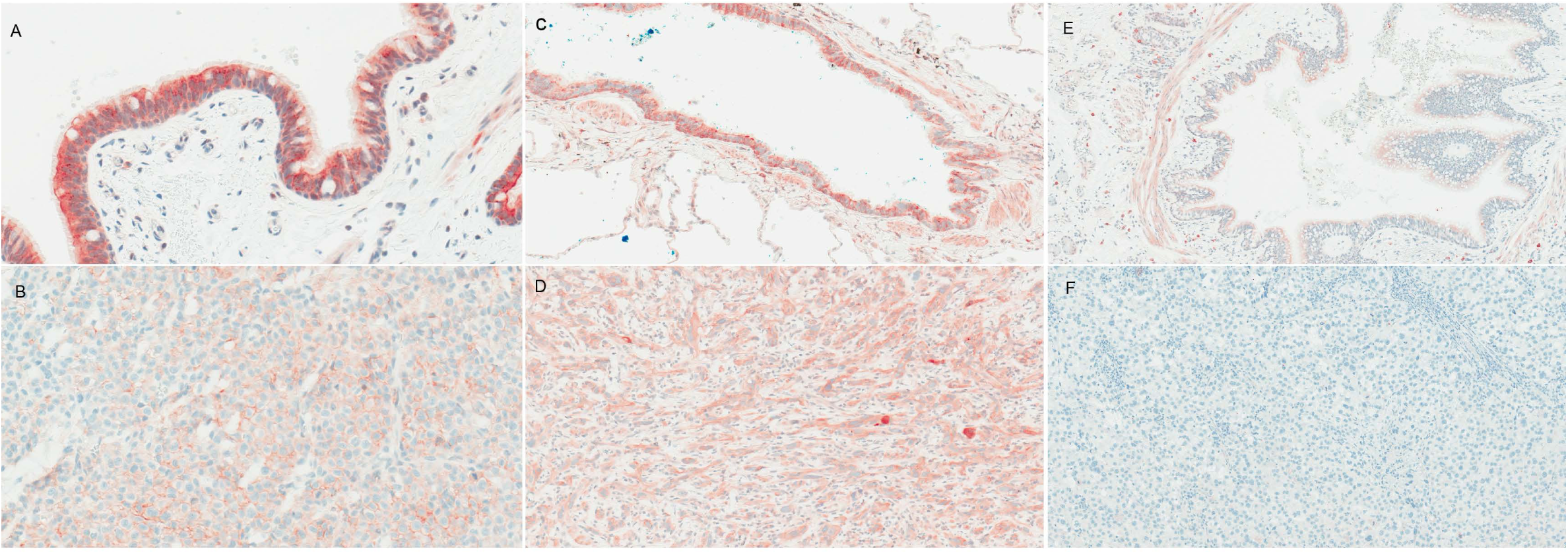
pβcat staining of lung tissue and melanoma metastasis from the same patient. Representative sections of pS33-βcat stained tissue from five cases of metastatic melanoma to lung at a 75x magnification. (**A, C, E)** staining of uninvolved respiratory epithelium from three tumors. (**B, D, F**) the corresponding metastatic melanomas from these cases.

**Figure 4.**
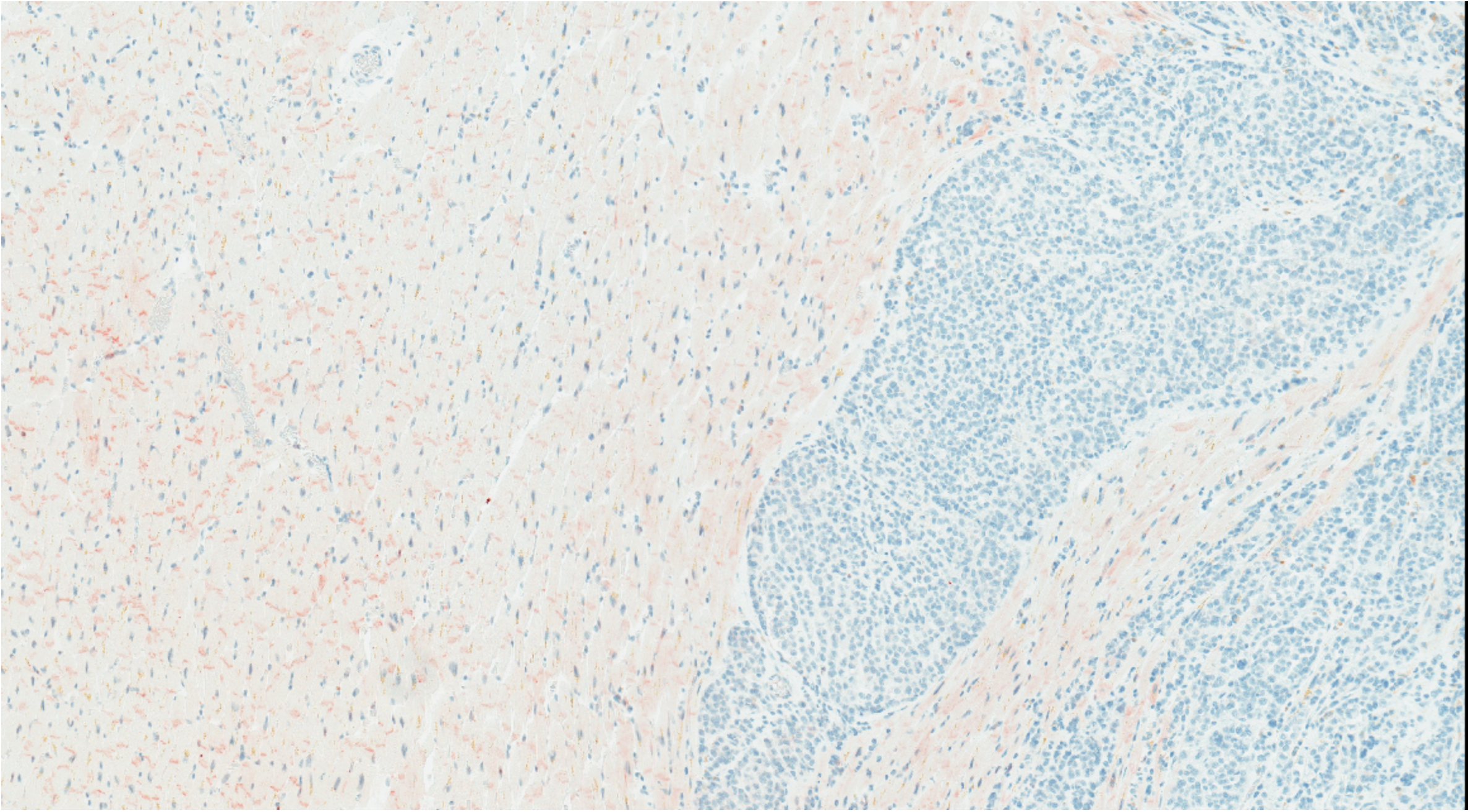
pβcat staining in heart tissue with melanoma metastasis. Representative section of pS33-βcat stained heart tissue from a patient with metastatic melanoma to heart at a 30x magnification. The uninvolved heart tissue is on the left side of the figure and has weak cytoplasmic staining with more intense staining of the intercalated discs. The metastatic melanoma on the right side of the figure has no significant staining.

**Figure 5.**
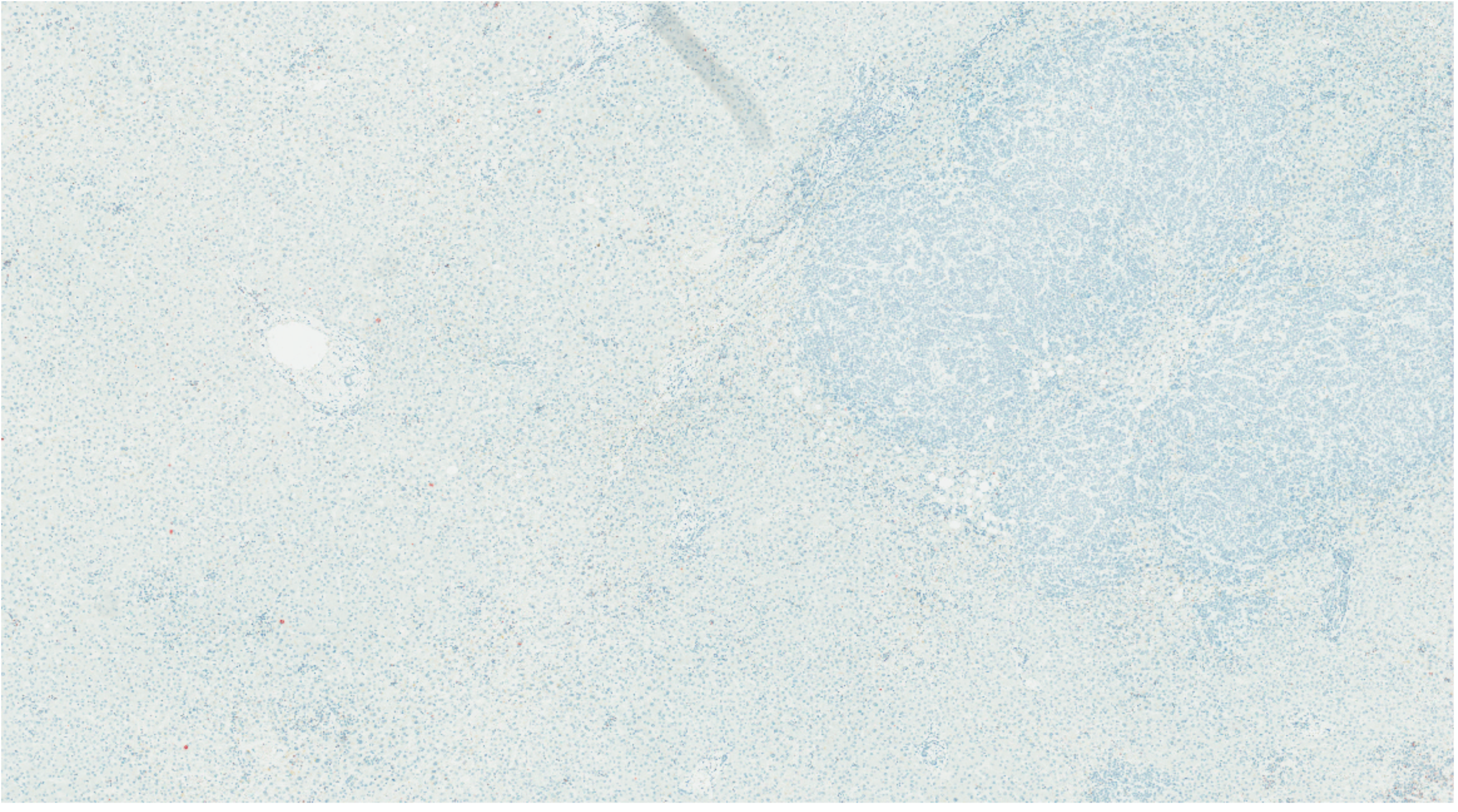
pβcat in liver tissue with melanoma metastasis. Representative section of pS33-βcat stained liver (20x magnification) containing a melanoma metastasis. Neither the uninvolved liver (left side of figure) nor the melanoma (right side of figure) had significant staining for pS33-βcat.

Phosphorylation of pS33 requires prior phosphorylation of S37 and T41 by GSK-3β^19^. We therefore hypothesized that an antibody to pS33-βcat would produce comparable staining to an antibody to pS33/37/T41-βcat. We stained an SLM-2 melanoma cell line with both antibodies (**Figure 6**). Indeed, the two antibodies produced comparable staining. For both antibodies, staining was largely cytoplasmic, with a minority of cells having combined nuclear/cytoplasmic staining. Nuclear staining was somewhat more evident with the pS33/37/T41-βcat antibody than the pS33-βcat antibody. One particularly striking finding was that mitotic cells showed much more intense staining with both pβcat antibodies than melanoma cells without mitotic figures.

**Figure 6.**
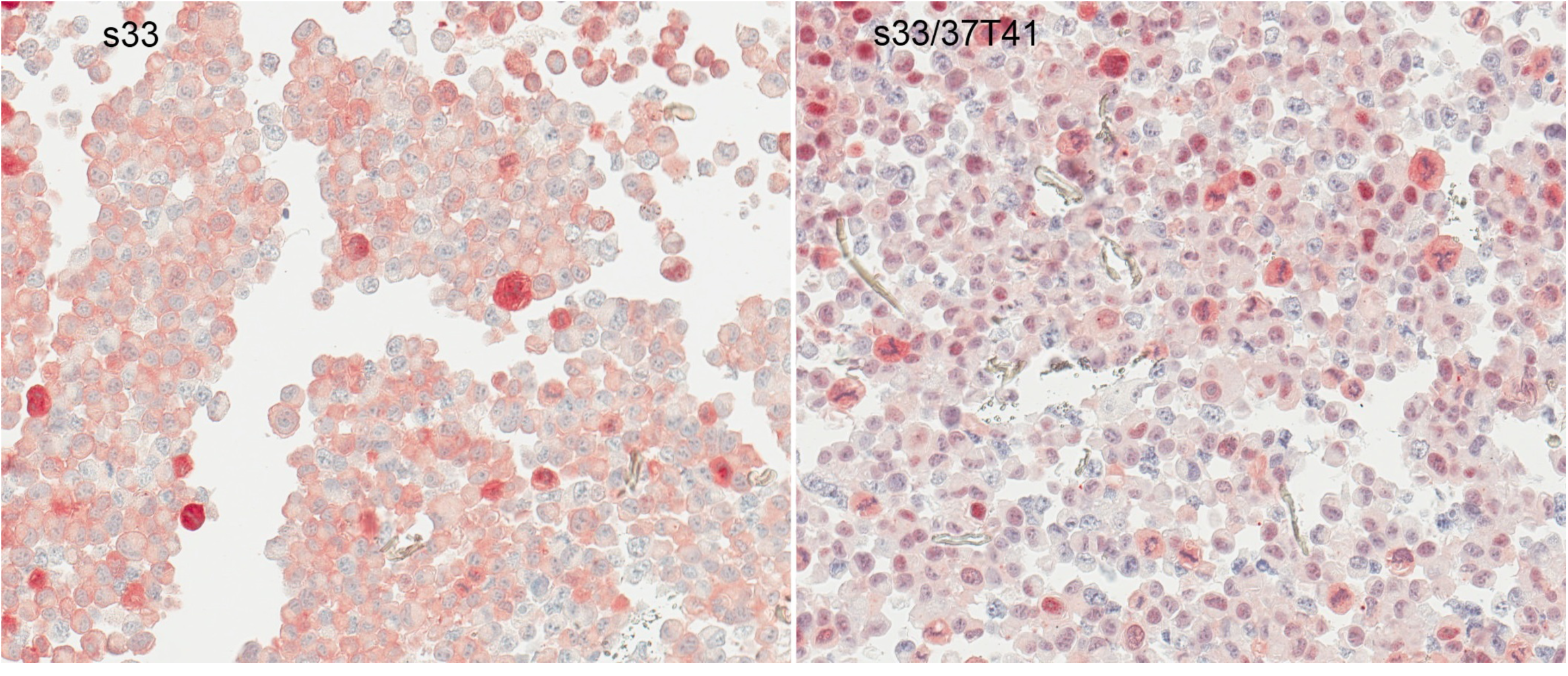
pβcat staining of melanoma cell block with two antibodies. A cell block of the SLM2 melanoma cell line was stained with antibodies to pS33-βcat (left panel) and pS33/37/T41-βcat (right panel). Staining for both antibodies was predominantly cytoplasmic but also included some nuclear staining with a slight predilection for nuclear staining with the pS33/37/T41-βcat antibody. Mitotic cells stained much more intensely than adjacent non-dividing cells.

## Discussion

We have found that melanoma metastases express pS33-βcat at varied levels and that the most common staining pattern in melanoma cells was cytoplasmic, both for the primary melanomas and for metastases. In contrast to expectation, there also was high expression of pS33-βcat in multiple normal tissues, including liver, placenta, spleen, kidney, lung, and heart. The validity of the antibody stain was confirmed by blocking with a specific phospho-β-catenin peptide. When evaluating lung and heart tissue containing melanoma metastases, the expression of pS33-βcat was higher in the surrounding normal tissue than in melanoma cells of the same specimen. A striking finding, however, was that the intensity of pS33-βcat staining in melanoma cells was greatest in actively dividing cells.

βcat serves an important function in embryogenesis and tissue development and is also an important downstream mediator of the development and progression of cancer. The N-terminus is of particular interest since this region contains phosphorylation sites that are critical for regulation of cellular levels of βcat that in turn affect the protein’s transcriptional activity. It has been suggested that pβcat expression may have diagnostic and/or prognostic value and specifically that nuclear staining of melanoma by an antibody to pS33-βcat, and the intensity of this staining might be useful as diagnostic markers to differentiate primary melanoma from metastatic melanoma and as prognostic markers in melanoma(14). In contrast to prior work(14), we found that cytoplasmic pS33-βcat staining was the most common staining pattern. In fact, exclusively nuclear staining only occurred in a single melanoma, and a complete lack of staining was also uncommon. Our finding of widespread cytoplasmic pS33-βcat staining in melanoma is much more in keeping with what might be expected if GSK-3β activity is maintained in the majority of melanomas. Kielhorn acknowledged that their finding was paradoxical, and suggested that it might be due to a defect in the proteasomal breakdown machinery in melanoma cells(14). However, while this would explain why pβcat might build up in the cells, it does not explain why the protein would be exclusively found in the nucleus. Our findings using both pS33-βcat antibodies are consistent with expectation that the most common location of pβcat is the cytoplasm.

Kielhorn also reported that more intense nuclear staining with the pS33/37/T41-βcat antibody was associated with metastatic melanoma(14). However, we found no significant differences between the intensity of either nuclear or cytoplasmic staining in the primary and metastatic melanomas. To evaluate whether pβcat expression differed between melanomas and normal tissues in vital organs, we selected tissue blocks containing metastatic melanoma and attached uninvolved tissue from the invaded organ to exclude tissue handling as a source of variability. Although the number of metastatic melanoma tissues tested in this study was limited, we found that cytoplasmic pβcat staining was less intense in the metastatic melanomas than in the uninvolved tissues in most cases. Although this could be due to a decrease in βcat phosphorylation in the metastatic cells, it seems unlikely since mutations in the N-terminus of βcat or in the components of the destruction complex such as APC are rare in melanoma(2, 5). Alternatively, these findings may also reflect an overall decrease in the level of βcat in melanoma, as previous studies have shown(18-20). This may be due to enhanced degradation of the protein as opposed to decreased gene expression since increased levels of mRNA have been found in metastatic melanoma tissues compared to primary melanoma(21). It is also possible that because melanoma cells are actively in cycle, pβcat is degraded more rapidly in melanoma cells than in normal tissues. βcat has been shown to accumulate in the cytoplasm during the cell cycle, with peak levels evident in the G2/M phase(22). Degradation of βcat, by phosphorylation, is needed for the cells to progress from the G2 phase through the M-phase(22). This correlates with our finding that mitotic figures in melanoma tissues show increased pS33-βcat staining intensity compared to cells not in the M phase of the cell cycle. Thus, the increased frequency of cell division in melanoma may account for an overall low level of pβcat while the increased levels observed in normal tissues may reflect steady state levels and a decreased rate of degradation owing to decreased frequency in cell division.

To our knowledge, the expression of pβcat in normal tissues has not been reported elsewhere. We found that normal heart and lung tissue both have high levels of pS33-βcat, with the most intense staining in the intercalated discs of myocytes and respiratory epithelial cells, respectively. βcat forms part of the adherens junction in intercalated discs. βcat has previously been reported to accumulate in the intercalated discs of myocytes in patients with hypertrophic cardiomyopathy(23). Furthermore, Maher et al have reported that pβcat and components of the destruction complex (axin, APC2, and GSK-3β) can localize to areas of cell-cell contact that are distinct from cadherin-catenin complexes at the cell membrane and facilitate phosphorylation of βcat, thereby leading to speculation about the possible role of βcat in cell fate during development and tissue morphogenesis(24).Thus, the high-level expression of pS33-βcat at the intercalated discs of the myocytes may be due to accumulation and phosphorylation of βcat at that site in normal tissues. Collectively, the results from our assessment of primary and metastatic melanomas as well as normal tissues suggest that pS33-βcat levels and localization are not suitable as either a diagnostic or a prognostic marker in melanoma.

One further interesting finding from our work was that pβcat staining was sharply increased in mitotic melanoma cells with antibodies both to pS33-βcat and to pS33/37/T41-βcat. This may be due to an increase in cytoplasmic βcat during mitosis, which is subsequently phosphorylated by GSK-3β leading to increased pβcat production. Indeed it has been shown that βcat and APC, a key component of the destruction complex, accumulates in the cytoplasm during the cell cycle, with the highest level of βcat being detected at the G2/M phase followed by a sharp decrease in βcat levels when cells enter the G1 phase(22). Additionally, the cells arrest in the G2 phase and undergo apoptosis if degradation of βcat is blocked by inhibiting GSK-3β, suggesting that phosphorylation of βcat at S33/37/T41 is necessary for cells to progress through the M phase. Thus, the strong pS33-βcat staining we observed in the mitotic melanoma cells is likely indicative of the imminent degradation of βcat in melanoma cells in this phase of the cell cycle. This finding may be useful in assessing whether pβcat could be targeted for cancer immunotherapy.

Overall, our results indicate that the pS33-βcat and pS33/37/T41-βcat antibodies produce comparable staining patterns in primary and metastatic melanoma. Cytoplasmic pβcat is most predominant in the tumor cells. We did not detect differences between pβcat expression between primary and metastatic melanomas; so, our data do not support use of pβcat antibodies for diagnostic or prognostic purposes in melanoma. Interestingly, we observed strong pβcat staining in mitotic melanoma cells. Given the results of other studies demonstrating that degradation of βcat is necessary for progression through the M phase, the phosphorylated peptide derived from βcat that was identified on melanoma cell lines may be an interesting target for immunotherapy for melanoma patients.

## Acknowledgements

This work was supported by the United States Public Health Service Grants R01CA134060 and R01CA190665 (VHE), T32AI007496 (JP), F31CA119954 (RCO), and T32GM007267 (RCO). We thank the University of Virginia Biorepository Tissue Resource Facility (supported by USPHS Cancer Center Support Grant P30 CA44579) for tissue sectioning and Donna H. Deacon for technical assistance.

